# A multimodal brain phantom for noninvasive neuromodulation

**DOI:** 10.1101/2025.10.13.682205

**Authors:** John LaRocco

## Abstract

Noninvasive neuromodulation enables brain stimulation without surgery but requires precise optimization of stimulation parameters to ensure efficacy and safety. Direct testing on human or animal subjects is costly, time intensive, and constrained by ethical and safety considerations. To address these challenges, a low-cost, versatile brain phantom was designed to emulate key biophysical properties across multiple neuromodulation modalities. The phantom was fabricated using ground beef, sodium alginate, and starch, and cast within a custom 3D printed mold. A multimodal test platform was created and validated by integrating established principles from low-intensity focused ultrasound (LIFU), transcranial direct current stimulation (tDCS), and thermal phantom design. Numerical simulations predicted a LIFU peak negative pressure of 0.631 MPa, closely matching the target Pr.3 value, with negligible temperature elevation (<0.01 °C). Consistent with prior reports, tDCS exposure did not induce lasting alterations in the phantom’s physical or electrical properties. The electrical conductivity was 0.11±0.02 S/m, reflecting water saturation within the phantom matrix; the thermal conductivity averaged 0.557 W/(mK), consistent with reported values for brain tissue analogs. This study primarily evaluated LIFU and tDCS performance; future work should extend characterization to additional modalities such as deep brain stimulation and transcranial magnetic stimulation. Further assessment of the phantom’s optical properties would also facilitate photobiomodulation and photoacoustic imaging studies. Overall, this inexpensive, easily fabricated phantom presents a practical and adaptable platform for multimodal neuromodulation research and parameter optimization.

## Introduction

### Overview

Noninvasive neuromodulation provides deep brain stimulation (DBS) without surgery but requires precise parameter optimization. Direct testing on human or animal subjects is costly, time consuming, and carries safety risks (LaRocco et al., 2024). To address these challenges, researchers employ phantoms—engineered materials such as gels, solids, and liquids—which mimic the electrical or physical properties of brain tissue (Mei et al., 2023; Linde et al., 2025; Saint-Martin & Avanaki, 2025). However, most phantoms are modality specific and cannot replicate the full range of tissue characteristics required for comparative studies. Given that many clinical and research environments utilize multiple neuromodulation systems, the development of a low-cost, versatile phantom modeling brain properties across modalities would significantly enhance the efficiency and reliability of parameter optimization.

## Background

### Multimodal stimulation

Brain phantoms for multimodal stimulation—employing neuromodulation methods such as electrical, magnetic, and ultrasound approaches—have evolved primarily in response to the need for realistic and experimentally controllable models for testing and optimizing brain stimulation techniques (Bystritsky et al., 2016; Gloeckner et al., 2021). Early phantom models were simplistic and lacked anatomical/electrical fidelity to human brain tissue, limiting their utility in studying complex neuromodulation effects (Allen et al., 2006). The advent of 3D printing and advanced conductive materials has facilitated considerable progress in recent years, enabling the construction of anatomically accurate brain phantoms which closely mimic the geometry and heterogeneous electrical conductivity of gray matter, white matter, and cerebrospinal fluid (Bystritsky et al., 2016; Choi et al., 2024). This allows for more precise modeling of the electric and magnetic fields induced by stimulation techniques such as transcranial magnetic stimulation (TMS), transcranial direct current stimulation (tDCS), and DBS (Bystritsky et al., 2016; Cho et al., 2023; Yeom et al., 2023; Aboalgasm et al., 2024; LaRocco et al., 2024).

This anatomical accuracy and the tunable electrical properties of these phantoms enable controlled experiments across multiple stimulation modalities simultaneously, a crucial advancement for multimodal neuromodulation research (Allen et al., 2006). A recently developed brain phantom integrates 3D printed shells with conductive polymers, enabling TMS, tDCS, and DBS studies in a manner closely replicating clinical conditions (Virginia Commonwealth University, 2025). Such phantoms facilitate parameter optimization prior to clinical application and can be individualized using patient-specific MRI data to tailor neuromodulation therapies (Cho et al., 2023; Yeom et al., 2023; Aboalgasm et al., 2024).

Parallel to hardware development, the concept of multimodal neuromodulation has gained traction for its potential synergistic effects, combining modalities such as electrical, magnetic, focused ultrasound, and optical stimulation to achieve enhanced spatial precision, specificity, and efficacy (Esmaeilpour et al., 2017; Yavari et al., 2018; Shiozawa et al., 2021). Multimodal phantoms are essential in validating these combinations, exploring interactions between stimulation modalities which may alter neuronal responsiveness and network dynamics (Sedley et al., 2019; Smith et al., 2025). For instance, the combination of ultrasound with electrical or magnetic stimulation can potentiate neural activation or modulation beyond that achieved by single modalities alone, an effect which phantom studies help elucidate under rigorously controlled conditions (Yang et al., 2022; Schilling et al., 2023; Weisz et al., 2023).

The progression of brain phantoms from rudimentary conductive models to anatomically faithful, multimodal-compatible constructs parallels advances in neuromodulation technology. These phantoms now underpin investigations into mechanisms of multimodal stimulation and optimization of therapeutic strategies targeting brain disorders, providing indispensable experimental platforms bridging computational modeling and clinical translation (Kraus et al., 2020; Cho et al., 2023; Yeom et al., 2023; Aboalgasm et al., 2024; Kraus et al., 2024).

### Electrical phantoms

One of the first electrical neuromodulation techniques, tDCS applies weak direct electrical currents across the scalp to modulate cortical excitability (Esmaeilpour et al., 2017; Yavari et al., 2018). Investigations into transcranial electrical stimulation began with Giovanni Aldini who applied galvanic currents for neurological and psychiatric conditions in the early 19th century. Modern controlled tDCS emerged primarily after the late 1990s, with foundational work demonstrating its ability to modulate cortical excitability in human and animal models (Jackson et al., 2016; Shiozawa et al., 2021).

Brain phantoms for tDCS are physical models developed to replicate the anatomical and electrical properties of the human head, facilitating direct measurement and validation of electrical current flow and electric field distributions during stimulation (Woods et al., 2016; Bikson et al., 2019; LaRocco et al., 2024). Due to the high reported variability of tDCS and other transcranial electrical stimulation techniques, these phantoms complement computational models by incorporating realistic tissue conductivity and complex anatomical geometries to better elucidate the spatial distribution of the applied currents within the brain (Polanía et al., 2019; Bikson et al., 2020).

The application of brain phantoms to tDCS research has evolved with technological advances in 3D printing and MRI-based anatomical modeling. Recently, high-fidelity 3D head phantom models have been fabricated from MRI-derived molds. These generally utilize tissue-mimicking materials such as agar to simulate scalp, skull, and brain compartments with embedded electrodes for voltage sensing at specific brain regions (Bikson et al., 2020; European Stroke Organisation, 2020). Such models allow for comparison of experimental measurements with physics-based simulations of electrical potentials and fields, supporting optimization of electrode configurations and stimulation protocols (Esmaeilpour et al., 2017; Yavari et al., 2018; LaRocco et al., 2024).

Studies using these phantoms have shown satisfactory agreement between measured voltages and computational predictions, demonstrating their utility in validating current flow models (Buch et al., 2021). For example, circular stimulation electrodes were found to better confine electric fields compared with rectangular ones, a key insight for improving focality in tDCS applications (Esmaeilpour et al., 2017; Yavari et al., 2018; Shiozawa et al., 2021; LaRocco et al., 2024). Brain phantom studies also highlight practical issues, such as electrode–skin contact impedances, which substantially affect current delivery and must be accounted for in modeling and clinical implementation (Shiozawa et al., 2021).

The history of brain phantoms for tDCS reflects a progression from rudimentary early electrical stimulation to sophisticated anatomically realistic physical models which serve as critical tools for validating and optimizing transcranial electrical stimulation techniques (Esmaeilpour et al., 2017; Yavari et al., 2018; Shiozawa et al., 2021). These phantoms continue to enhance understanding of the biophysics of tDCS and support translation into therapeutic and research applications (Buch et al., 2021; Shiozawa et al., 2021).

### Acoustic phantoms

The development of brain phantoms specifically used for low-intensity tFUS is closely tied to that of brain phantoms for ultrasound propagation and neuromodulation research, with emphasis on replicating skull and tissue properties relevant to ultrasound delivery (Zhang et al., 2021; Zhao et al., 2023). Early brain phantoms relevant to focused ultrasound research were designed primarily to mimic acoustic propagation characteristics through the skull and soft tissues to facilitate applications such as thermal ablation and noninvasive neuromodulation (Focused Ultrasound Foundation, 2025). Designs emerged to overcome challenges posed by the skull’s high attenuation and scattering of ultrasound which affect the precision and focality of energy delivery (Legon et al., 2014; Kuzmina & Bansal, 2023).

The rise of low-intensity focused ultrasound (LIFU) as a neuromodulation technique in the early 21st century intensified the need for brain phantoms simulating human cranial anatomy— including heterogeneous skull bone layers and brain tissue acoustic properties—allowing safe parametric evaluation of transcranial ultrasound targeting, intensity modulation, and dosimetry without the ethical or safety concerns associated with in vivo testing (Legon et al., 2014; Kuzmina & Bansal, 2023). Phantoms were developed to mimic both mechanical and acoustic tissue parameters such as speed of sound, density, attenuation, and acoustic impedance, critical for modeling ultrasound transmission and focal spot formation under low-intensity neuromodulation regimes (typically <50 W/cm² spatial peak temporal average intensity) (Bystritsky et al., 2016; Brainbox Neuro, 2019; Basso et al., 2024; Choi et al., 2024).

Technically advanced phantoms for low-intensity tFUS incorporate layered materials representing the scalp, skull cortical and trabecular bones, and brain parenchyma. These are often fabricated with polymers, hydrogels, or composite materials embedded with scatterers to emulate the ultrasound beam refraction, reflection, and absorption effects observed in human subjects. This has enabled quantitative analysis of beam focusing quality, pressure fields, and safety limits in neurological targets such as the motor cortex, thalamus, and hippocampus, where low-intensity pulsed ultrasound is used to modulate neural activity noninvasively with millimeter spatial precision (Bystritsky et al., 2016; Basso et al., 2024; Choi et al., 2024).

Some state-of-the-art phantom models are coupled with imaging modalities (e.g., MRI, CT) or embedded sensors to enable real-time monitoring and calibration of ultrasound exposure, critical for the translation of tFUS neuromodulation protocols to clinical applications. Such phantoms provide an indispensable testbed for optimizing ultrasound parameters, verifying neuromodulation effects, and ensuring reproducibility in preclinical and human studies (Fini & Tyler, 2020; Basso et al., 2024).

The evolution of brain phantoms for low-intensity transcranial focused ultrasound stimulation reflects a transition from early models emphasizing acoustic transmission for therapeutic ablation to sophisticated models replicating the heterogeneous and layered nature of the human head to support precise, safe, and targeted neuromodulation under low-intensity conditions (Bystritsky et al., 2016; Choi et al., 2024). These phantoms present foundational tools to characterize ultrasound propagation and focal stimulation effects noninvasively and in controlled laboratory settings, facilitating advances toward clinical neuromodulation harnessing the unique capabilities of low-intensity tFUS (Bystritsky et al., 2016; Basso et al., 2024; Choi et al., 2024).

### Thermal phantoms

The use of brain phantoms in heat transfer research originated from the need to investigate the thermal behavior of brain tissue under various therapeutic/diagnostic conditions (Packett et al., 2017). Early efforts in this domain aimed to replicate the brain’s thermal properties and response to blood perfusion and external cooling/heating (Xu et al., 2010; Cho et al., 2023). Some early phantoms utilized materials such as superabsorbent hydrogels to mimic the thermal conductivity, specific heat, and perfusion characteristics of cerebral tissue (Zhang et al., 2018). This enabled studies on heat transfer mechanisms during cooling therapies targeted at neurological conditions such as epilepsy and concussions (Cho et al., 2023; Yeom et al., 2023; Aboalgasm et al., 2024).

The field evolved to include anatomically accurate phantoms created using advanced fabrication techniques such as 3D printing, enabling more realistic simulation of brain geometry, including the differentiation between gray and white matter (Li et al., 2024). These phantoms allow for detailed analyses of heat transfer phenomena influenced by structural and material heterogeneity in the brain tissue and surrounding skull (Xu et al., 2010; LaRocco et al., 2024). The integration of perfusion effects, representing cerebral blood flow within phantoms, has become critical in accurately simulating thermal transport, as blood perfusion significantly influences heat dissipation in living brain tissue (Cho, 2023). Mathematical models, particularly those based on the Pennes bioheat equation, have been extensively employed to guide phantom design and verify experimental results (Farahani, 2019; Kainz et al., 2019; Aboalgasm et al., 2024).

Recent advancements include brain tissue/skull phantoms tailored to the study of focused ultrasound heating and thermal therapies, addressing the complex interactions of ultrasound with heterogeneous brain/skull materials (Aboalgasm et al., 2024). These phantoms facilitate the evaluation of energy deposition and heat diffusion relevant for clinical applications of thermal ablation and neurotherapeutics (Farahani, 2019; Kainz et al., 2019; Xu et al., 2019).

Brain phantoms for heat transfer have progressed from simple material analogues to sophisticated anatomically and physiologically representative models combining experimental phantom construction with bioheat modeling to support the investigation of therapeutic heating and cooling effects in the brain (Cho et al., 2023; Yeom et al., 2023; Aboalgasm et al., 2024).

## Methods

### Overview

Prior brain phantoms have comprised solids, gels, liquids, and other materials (Mei et al., 2023; Linde et al., 2025; Saint-Martin & Avanaki, 2025). Materials within the phantom can be customized for specific properties such as acoustic impedance and electrical resistance. Prior work described a low-cost phantom for electrical properties made from ground beef (LaRocco et al., 2024). Cornstarch has been used in acoustic phantoms to adjust acoustic impedance; gel binders such as sodium alginate provide structural stability (Hellerbach et al., 2013; Huo et al., 2015; Kilian et al., 2022). Without a circulatory system or active cooling, many phantoms have no facility for temperature reduction. As such, simple phantoms possess a lower thermal conductivity than living tissue (Islam et al., 2021). Testing brain phantoms alongside LIFU stimulation and AI could enable safer optimization of neuromodulation parameters.

### Neuromodulation modalities

Three types of neuromodulation were investigated: LIFU, tDCS, and laser. Each modality characterized a different aspect of the phantom: acoustic, electrical, and thermal. LIFU was conducted using the same parameters as prior work (Jenkins et al., 2025); the tDCS method used the same device as a prior phantom study (LaRocco et al., 2024); and the third method utilized a 447 nm blue laser (MDL-XS-447), using a 2.4-W beam. As the properties of the brain and phantom material were known, the values for acoustic impedance, electrical resistance, and thermal conductivity could be directly compared with previously reported values.

### Phantom model

As shown in Fig 1, the phantom consisted of meat, a plastic shell, and a gel mixture. A clam-shell plastic case was based on the typical shape and volume of a human brain at 1,158 cm³, held together with screws and clamps (Rushton & Ankney, 2009; Zhang et al., 2017). The plastic shell was 3D printed with polylactic acid with 20% infill and weighed 425 g. Polylactic acid has previously been used as an analog of the human skull (Zhang et al., 2017; Antoniou & Damianou, 2024). Approximately 1.06 kg of ground beef was used as the base of the phantom (LaRocco et al., 2024). A total of 17 g of cornstarch was added to adjust acoustic impedance (Islam et al., 2021). To improve structural stability, 14 g of distilled water and 7 g of sodium alginate were combined and applied to the container (Hellerbach et al., 2013; Huo et al., 2015; Kilian et al., 2022). The mixture was manually shaken for 2 minutes, drained of water, and cooled in a controlled climate of 4 °C for 12 hours prior to experimentation.

**Fig 1:**
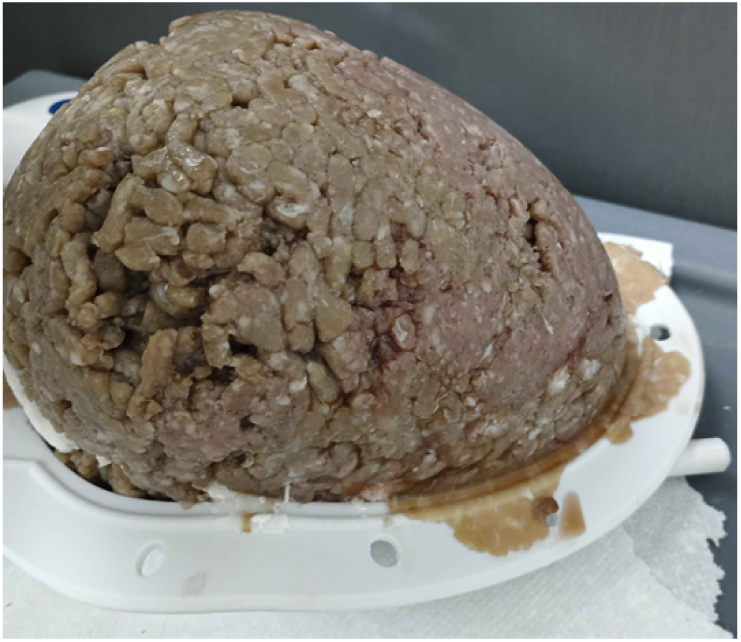
Side view of the brain phantom in the plastic shell.

### Values from the literature

Modeling the human brain requires a phantom with comparable mechanical, acoustic, electrical, and thermal characteristics. In mechanical scenarios, the Young’s modulus of the brain averages 8 kPa (range, 5.5–14 kPa) under separate experimental conditions (Bennion et al., 2022). Inclusion of the skull and skin adds another layer, highly relevant for acoustic impedance due to the change in medium. Prior work reported the acoustic impedance for 650 kHz at 1.7 MRayl (Tang et al., 2025). The electrical conductivity of the skin and brain is relevant for electrical characteristics, as current flows along the path of least resistance. A conductivity of 0.44 S/m was reported for the human brain, with aqueous cerebrospinal fluid comparable to water (McCann et al., 2019). Thermal conductivity is relevant for a heat source applied to the brain, to direct the speed of heat accumulation in one part of the brain. Based on specific frequencies, temperature, and power, these coefficients vary significantly by the specific neuromodulation parameters. In the brain, thermal conductivity at the physiological temperature range was estimated at 0.56 W/(mK) (Mohammadi et al., 2021). In all cases, the “dosage of power” (Dp) was the product of the energy (E) and time (t), as shown in Eq 1.

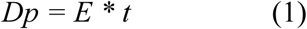

The dosages for each modality are detailed in Table 1.

**Table 1:** Energy and power delivered per stimulation dose, including LIFU, tDCS, and laser.

| Method | Power (W) | Time (s) | Dose (J) |
| --- | --- | --- | --- |
| LIFU | 0.716 | 30 | 21.48 |
| tDCS | 0.018 | 1,200 | 21.6 |
| Laser | 2.4 | 10 | 24 |

### LIFU neuromodulation

As with prior work targeting the left amygdala, the MR-guided treatment was performed using the BrainSonix Pulsar 1002 LIFU pulsation system equipped with a 65-mm transducer MRI compatible up to 3 T (Schafer et al., 2020; Jenkins et al., 2025). As with human participants, the transducer was secured with elastic straps and 5°-angled gel pads to minimize dispersion and ensure effective acoustic coupling (Fonzo et al., 2021). In its default configuration, the sonication beam had a target depth of 65 mm.

Once properly positioned, as shown in Fig 2 and Fig 3, the LIFU transducer delivered 30-second trains of 650 kHz sonication at a pulse repetition frequency of 10 Hz, interspersed with 30-second rest periods. This protocol was repeated for a total of 10 cycles over 10 minutes, with a peak sonication intensity of 720 mW/cm² (Radjenovic et al., 2022; Spivak et al., 2022b). The pulse width was 5 ms; the duty cycle was 5%. The mechanical index of the transducer was 1.7, below the limit of 1.9. These parameters are consistent with those used in other studies investigating amygdala modulation via LIFU stimulation (Fonzo et al., 2021; Philip & Arulpragasam, 2023).

**Fig 2:**
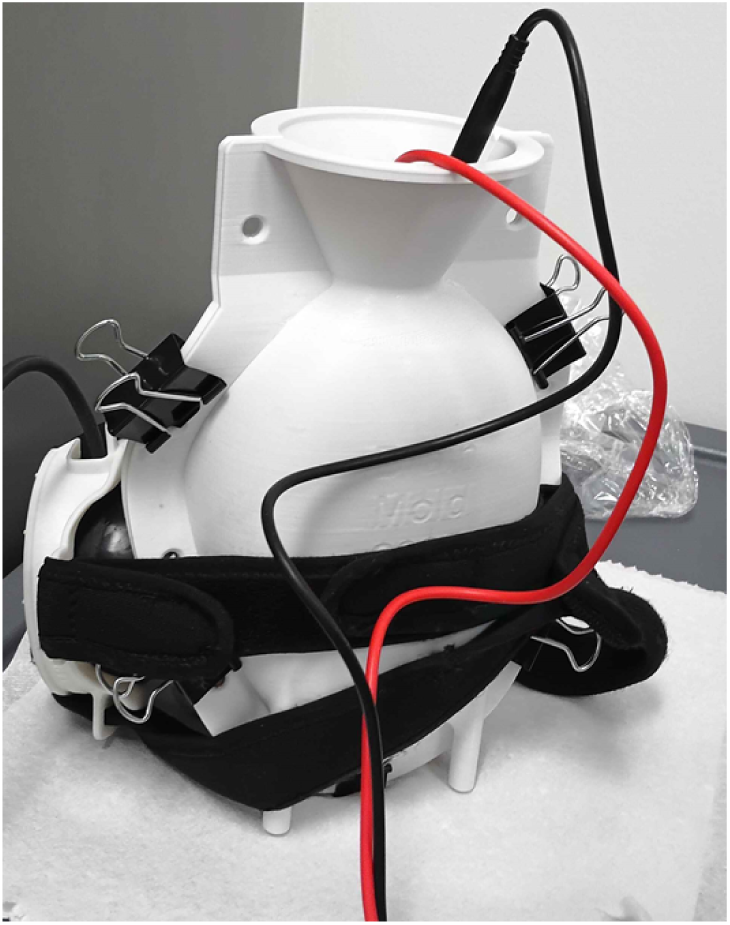
Positioning the transducer on the phantom casing.

**Fig 3:**
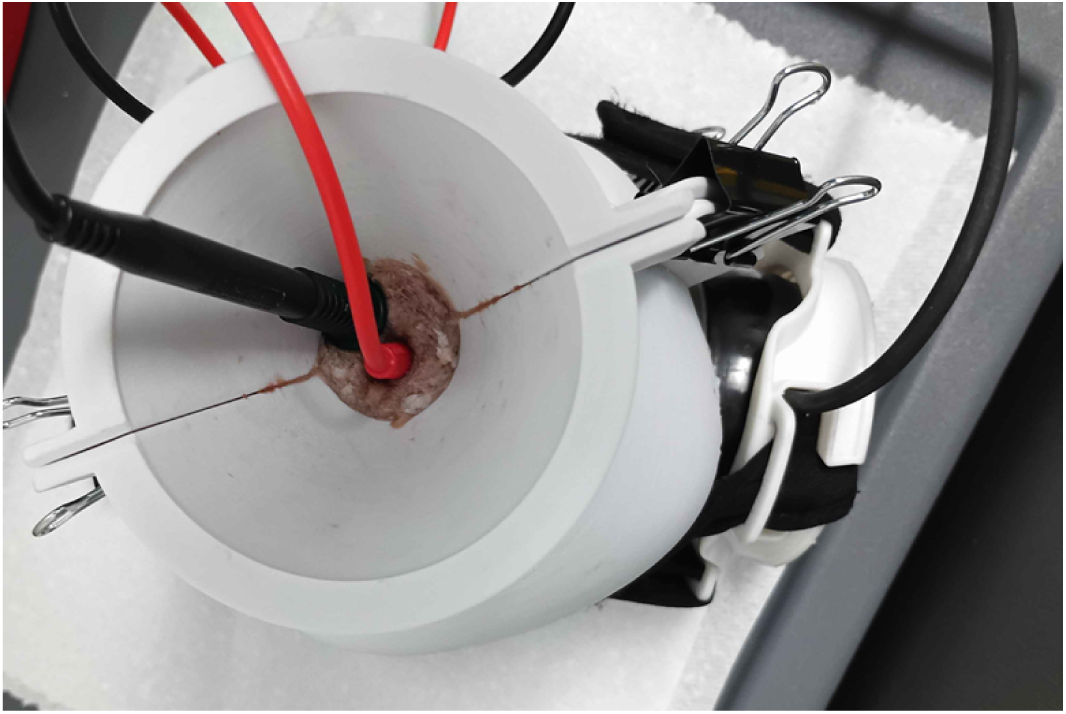
Top view of transducer positioning on the phantom.

### Temperature and pressure changes

A simulation was constructed to estimate temperature and pressure changes. Based on prior experimental values, the model was a Gaussian beam with a depth of 65 mm, including a skull thickness of 3 mm. Skull attenuation was assumed to be 2 dB/mm. As with similar studies, cerebrospinal fluid was modeled as water due to its aqueous composition. The speed of sound in the brain was modeled as 1,540 m/s (Pichardo et al., 2010; Wang et al., 2014). The density (p) was modeled as 1,000 kg/m³. The specific heat (Cp) of the tissue used was 4,200 J/kg*K. Following from the ISPTA.3 of 716 mW/cm², the specific volumetric heating rate (Q) was 7,160 W/m². Attenuation for the system was 0.75 B/(MHz^y cm), and a power law exponent of 1.1 was adjusted for the skull and brain (Sukstanskii & Yablonskiy, 2006; Pichardo et al., 2010; Wang et al., 2014). The volumetric heat (Qvol) was modeled as a relationship between (Q) and a predefined coefficient.

Temperature was modeled using the bioheat equation (Sukstanskii & Yablonskiy, 2006; Pichardo et al., 2010; Wang et al., 2014) in Eq 2.

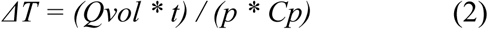

Following simulation in k-Wave software (Treeby et al., 2010), the results were compared with experimental measurements.

### Electrical properties

Ground beef is a widely used electrical brain phantom, owing to its similar composition of amino acids, salts, lipids, and proteins (LaRocco et al., 2024). The total composition of non-ground beef materials (including sodium alginate, starch, and any residual water), was less than 3.9% by weight; therefore, only the resistance of ground beef was used. At room temperature, the conductivity of ground beef depends on its degree of leanness or fattiness, ranging from 0.001–0.1 S/m. Conductivity can be higher, such as 1.6 S/m, where the water content is lower (Bozkurt & Icier, 2010). Using prior and estimated values for the size of the phantom and the presence of sodium alginate and residual water with resistivity testing, a resistance of 1 K-ohms was determined (Bozkurt & Icier, 2010; LaRocco et al., 2024).

### Electrode placement

Mechanical, thermal, and electrical stimulation can induce current flows in phantom models. As with prior work, multimeter electrodes were adapted and utilized for measurements (LaRocco et al., 2024). The voltage difference was measured between the surface and center of the phantom. As shown in Fig 4, one probe was placed within 1 cm of the surface and the other 10 cm into the center. Heat, motion, and vibration can induce electrical changes due to ionic salts, even in ground beef (LaRocco et al., 2024); however, the signal is often diminished and faint compared with living organisms.

**Fig 4:**
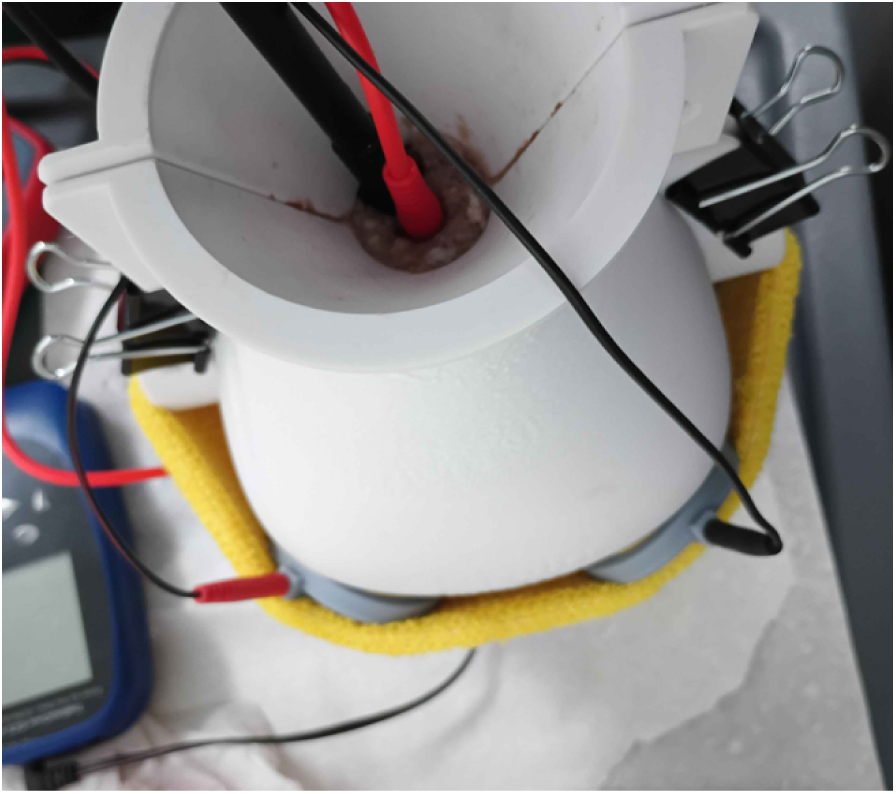
Positioning tDCS electrodes on the phantom.

### Electrical stimulation

Electrical stimulation was conducted using a commercially available tDCS device, a BrainDriver v2.1 (TheBrainDriver, Chicago, IL, USA). The BrainDriver delivered a constant DC voltage of 9 V, using a pair of sponge electrodes 2 cm in diameter. As with previous studies exploring dosages in brain phantoms, a current of 2 mA was applied for 20 minutes (LaRocco et al., 2024). Electrodes were positioned 10 cm apart on the front of the phantom, mimicking common tDCS montages (LaRocco et al., 2024). Electrical measurements were noted before and after stimulation. Measurements were used to calculate induced changes, allowing for the calculation of conductivity. In Eq 3, the power deposited (Pd) was the product of the voltage (V) and current (I). Using calculated and observed values, the tDCS device would be capable of depositing 0.018 W.

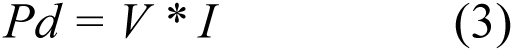

The total energy (E) the device was capable of depositing is shown in Eq 4, the product of power deposited (Pd) and time (t). Inputting earlier values, (E) was 21.6 J, as in prior work (LaRocco et al., 2024).

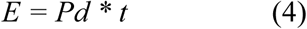

### Thermal stimulation

As shown in Fig 5, one half of the clamshell casing was removed to ensure that no energy was lost from the laser, and to prevent physical damage to the shell. Thermal stimulation utilized a 450 cm laser to deliver 2.4 W of thermal energy for 10 seconds. The laser required a warm-up time of approximately 3 seconds; thus, each position was stimulated for a total of 13 seconds. The laser was deactivated if smoke was detected, regardless of whether the total time had elapsed. The laser was positioned at intervals of 15° at four separate positions, roughly paralleling electrodes CZ, F7, T3, and T5 in the 10–20 international system for electroencephalography (LaRocco et al., 2024). The laser system circuit is illustrated in Fig 6. Using a contactless infrared thermometer, the temperature was measured at each location before and after stimulation, as well as at the top (near electrode position CZ) after each stimulation dose.

**Fig 5:**
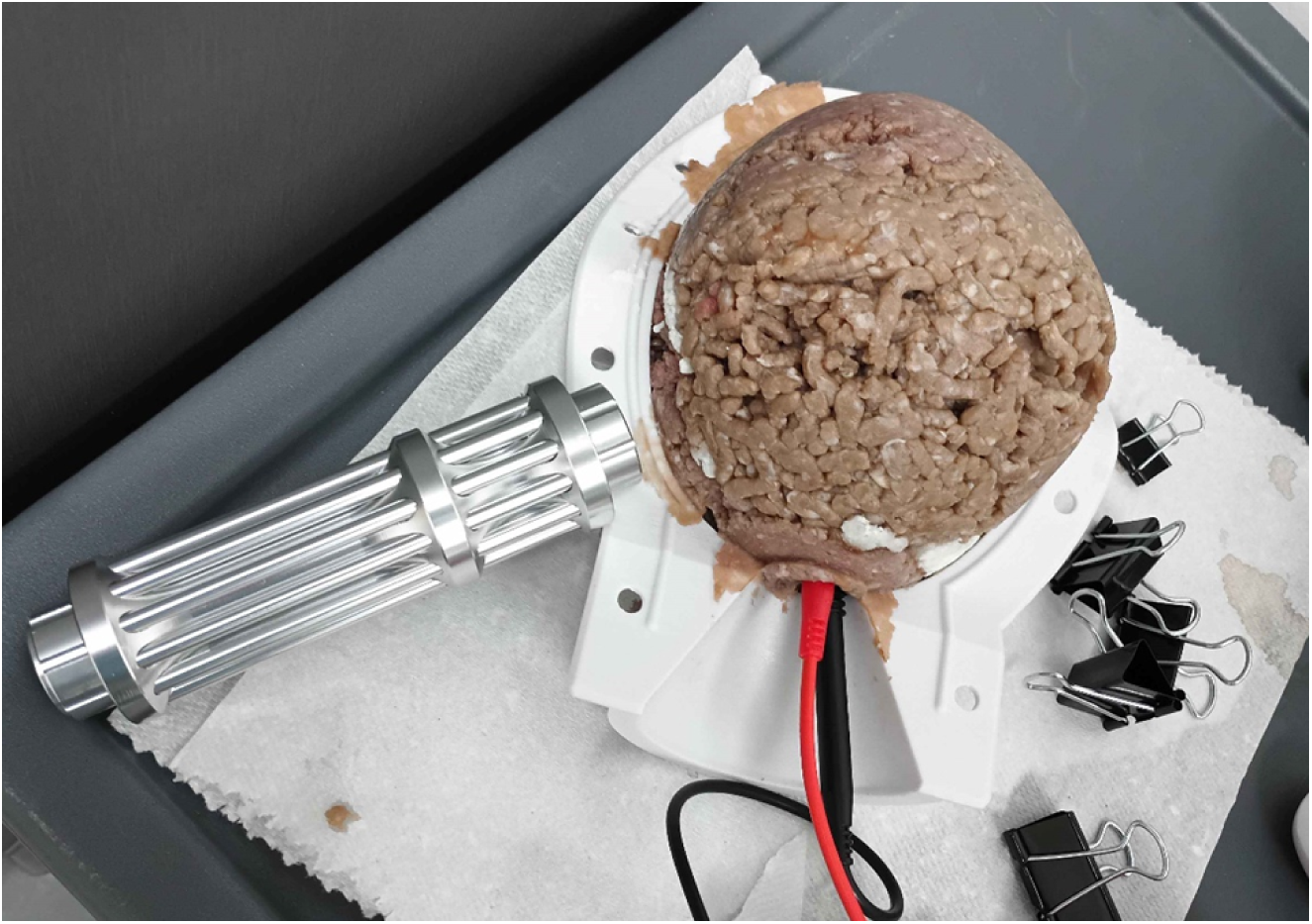
Laser positioning on the phantom.

**Fig 6:**
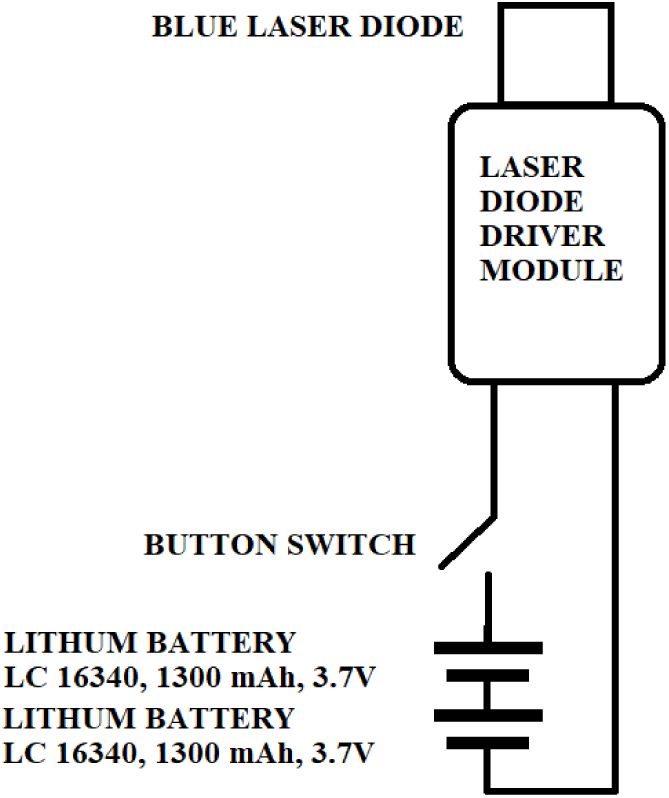
Laser power delivery circuit.

### Phantom estimates

The mechanical, acoustic, electrical, and thermal properties were estimated based on the composition of the phantom. Ground beef has a range of values for Young’s modulus far higher than the human brain, estimated at approximately 150 kPa (Kowal et al., 2015). For acoustic impedance, the combination of the casing and meat was assumed to approximate the skull and brain, with a 1.7 MRayl (Tang et al., 2025). Without water, electrical conductivity matched biological matter up to 1.6 S/m, and as low as 0.1 S/m (Bozkurt & Icier, 2010). With water, the conductivity dropped to a fraction. Due to the short distances between electrodes, a median resistance of 1 k-Ohms was assumed. It was assumed that the small additions to the ground beef would not significantly change the thermal conductivity of 0.56 W/(mK) (Mohammadi et al., 2021). These properties are compared with the brain in Table 2.

**Table 2:** Comparing physical properties of the phantom with values from the literature.

| Property | Units | Literature | Phantom |
| --- | --- | --- | --- |
| Acoustic Imp. | MRayl | 1.7 | 1.7 |
| Young's Mod. | kPa | 8 | 150 |
| Electrical Cond. | S/m | 0.44 | 0.1 |
| Thermal Cond. | W/(mK) | 0.56 | 0.56 |

### Data analysis

The phantom enabled direct measurement of thermal, electrical, and mechanical heating; however, the experiment was constrained in certain regards. Total immersion in water could compromise the phantom’s physical consistency; therefore, a hydrophone was unsuitable for the explored configuration. External thermal measurements and internal electrical activity were used as the primary observations. The electrode placement was fixed, but positioned in the center of the phantom to detect any electrophysical measurements in the deepest part (LaRocco et al., 2024). Three temperature and voltage measurements were noted, sampled at 10 Hz each. Adjusting for residuals, results before and after each stimulus were compared using a paired t-test with 95% confidence intervals in Python v3 (Amemaet, 2021). The temperature shift of the phantom toward room temperature was another potential complication; the phantom was therefore left at room temperature for 1 hour prior to commencement of the tests. Tests were conducted with the lowest power systems first, beginning with LIFU (acoustic), followed by tDCS (electrical), and concluding with laser (thermal).

### Experimental hypotheses

Due to the low amount of energy from LIFU, it was hypothesized that little difference would be detected. Other hypotheses were that the low amount of energy from thermal stimulation would translate to little observable difference in temperature; that electrical stimulation would raise electrical activity, and potentially, temperature with it; and that the laser would result in the highest amount of energy transfer, simply due to the overwhelming amount of energy contained. A significant response to thermal stimulation, and possibly electrical, indicates a closer model to the brain (LaRocco et al., 2024).

## Results

### LIFU simulation

Using a k-Wave simulation shown in Fig 7, the absolute average peak negative pressure was 0.631 MPa, slightly different from the stated Pr.3 value, 0.628 MPa.

**Fig 7:**
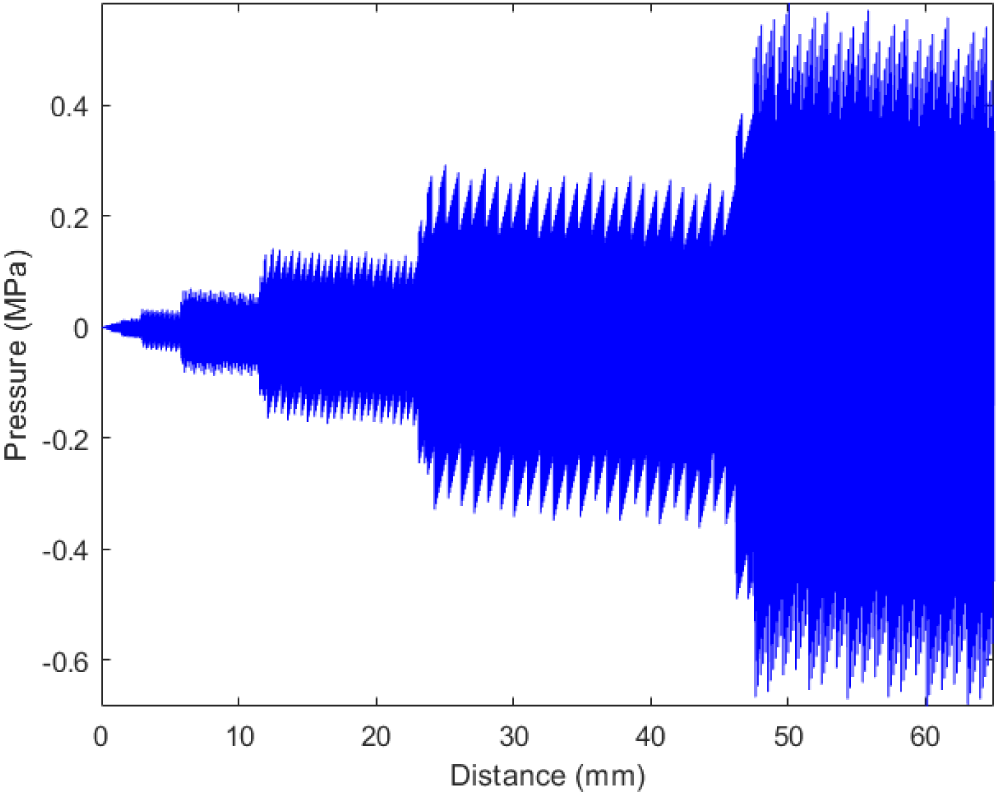
Pressure over distance from low-intensity focused ultrasound simulation.

Due to cerebrospinal fluid increasing the thermal mass and heat absorption capacity, the simulated change in temperature was 0.01 °C.

During LIFU testing, no significant changes in temperature were detected before and after stimulation. These findings are illustrated in Fig 8.

**Fig 8:**
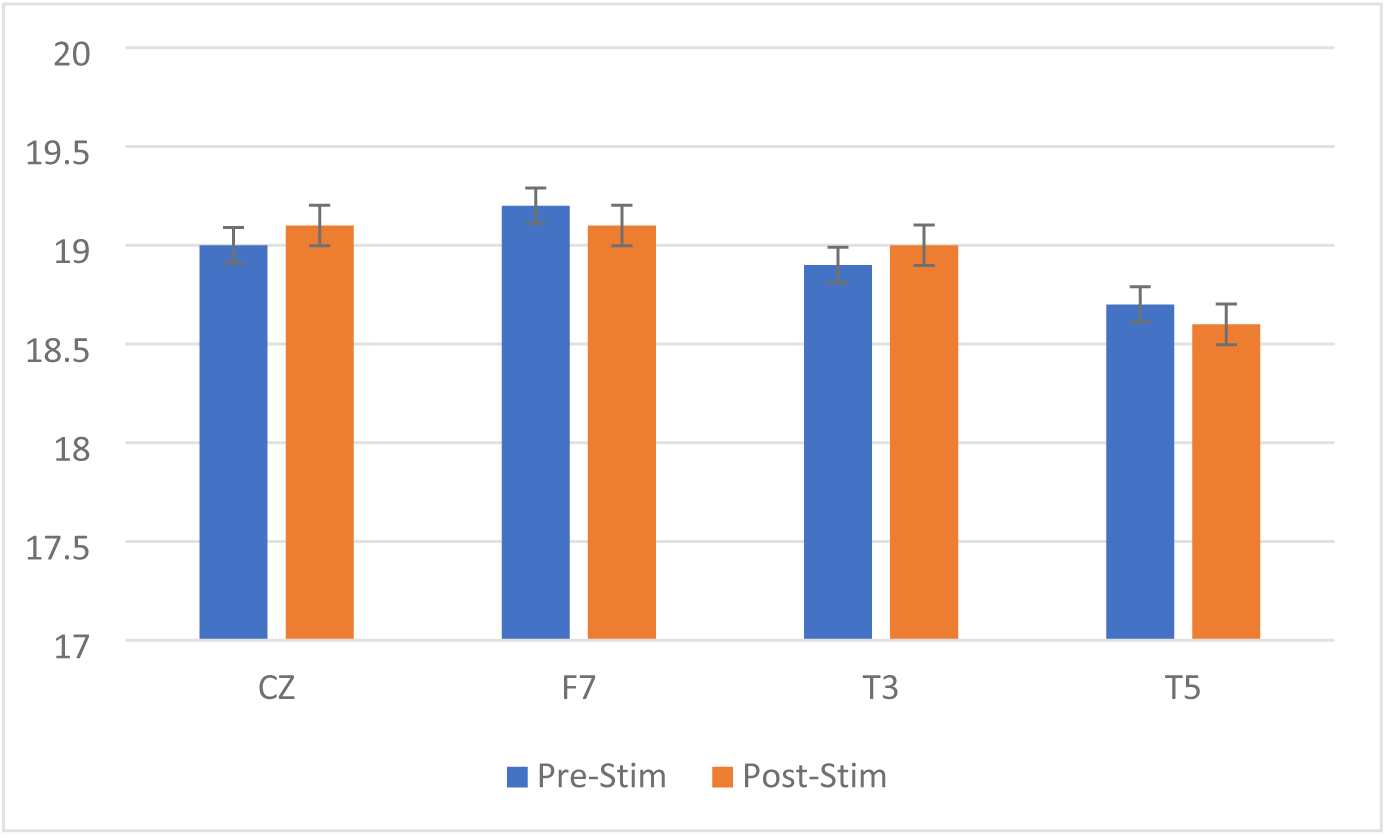
Comparison of temperatures (°C) before and after low-intensity focused ultrasound stimulation.

Table 3 compares experimental temperature measurements, showing no significant differences at each location following LIFU stimulation. Differences are within simulated parameters.

**Table 3:**
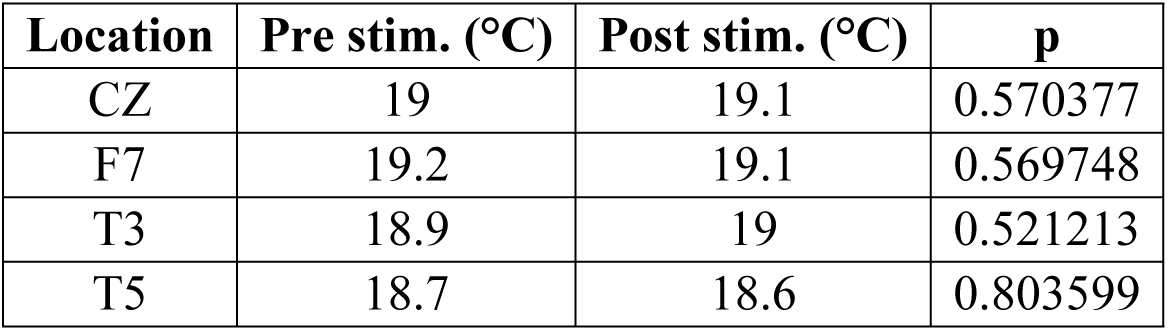
Temperatures before and after low-intensity focused ultrasound stimulation.

| Location | Pre stim. (°C) | Post stim. (°C) | p |
| --- | --- | --- | --- |
| CZ | 19 | 19.1 | 0.570377 |
| F7 | 19.2 | 19.1 | 0.569748 |
| T3 | 18.9 | 19 | 0.521213 |
| T5 | 18.7 | 18.6 | 0.803599 |

### Electrical measurements

Electrical changes were observed in each stimulation modality (Table 4). While some voltage changes were observed during LIFU, these may have been due to mechanically induced compression cycles on electrodes from active LIFU. The electrical conductivity averaged 0.11±0.02 S/m.

**Table 4:** Voltages before and after low-intensity focused ultrasound stimulation.

| Location | Pre stim. (mV) | Post stim. (mV) | p |
| --- | --- | --- | --- |
| CZ | 0.1 | 0.09 | 0.531805 |
| F7 | 0.12 | 0.11 | 0.767853 |
| T3 | 0.05 | 0.06 | 0.795167 |
| T5 | 0.09 | 0.1 | 0.424922 |

The tDCS was not sufficient to induce significant electrochemical or physiological changes, as 21.6 J is reported as a low, safe threshold (LaRocco et al., 2024).

### Thermal observations

No statistically significant differences were observed in average temperatures before and after stimulation. While the area directly stimulated by the laser increased rapidly, smoke rose within a few seconds. Local temperature changes dissipated rapidly. The presence of water, while slight, may have increased thermal absorption; however, acclimation to room temperature is another potential explanation. These results are detailed in Table 5 and Fig 9.

**Fig 9:**
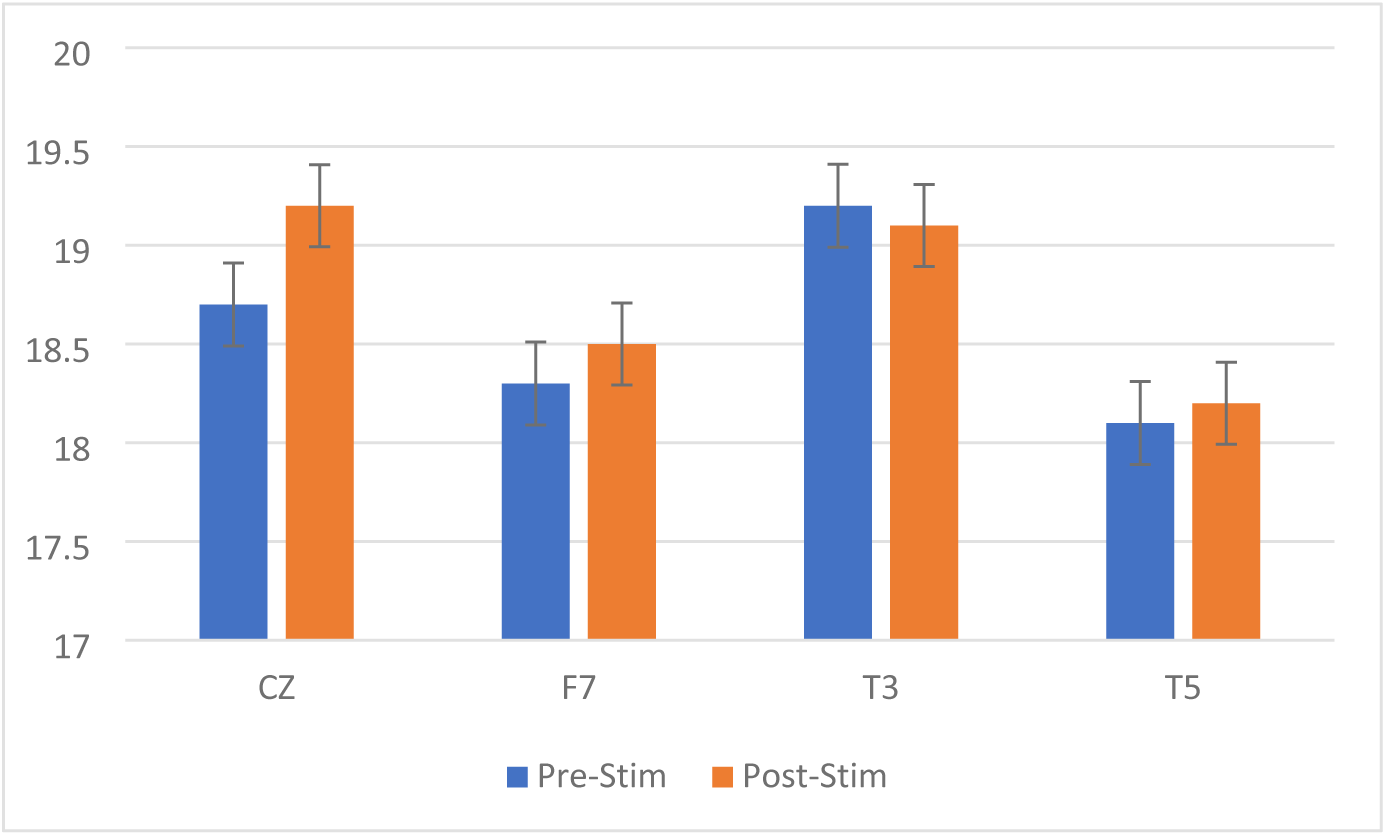
Comparison of pre- and post-laser stimulation temperatures (°C).

**Table 5:** Temperatures before and after laser stimulation.

| Location | Pre stim. (°C) | Post stim. (°C) | p |
| --- | --- | --- | --- |
| CZ | 18.7 | 19.2 | 0.338 |
| F7 | 18.3 | 18.5 | 0.434 |
| T3 | 19.2 | 19.1 | 0.768 |
| T5 | 18.1 | 18.2 | 0.741 |

The measured thermal conductivity was 0.557 W/(mK), comparable to the literature (Mohammadi et al., 2021).

## Discussion

### Overview

Multimodal neuromodulation integrates multiple forms of energy delivery, necessitating a brain phantom capable of replicating relevant physical and electrical properties (Bystritsky et al., 2016; Gloeckner et al., 2021). The developed phantom was evaluated under acoustic, electrical, and thermal stimulation to assess its multimodal response characteristics. The observed changes in temperature, electrical conductivity, and thermal conductivity were consistent with values reported in previous studies and simulation data. Simulations estimated a LIFU peak negative pressure of 0.631 MPa, closely matching the specified Pr.3, and producing a temperature rise of less than 0.01 °C; tDCS did not induce lasting alterations, consistent with previous findings (LaRocco et al., 2024). The electrical conductivity was 0.11±0.02 S/m, which remained relatively low due to water saturation within the phantom matrix (Mei et al., 2023; Linde et al., 2025; Saint-Martin & Avanaki, 2025). Minor electrical variations were observed during active LIFU application, likely attributable to transient electrode exposure or mechanical effects (LaRocco et al., 2024). Thermal stimulation resulted in rapid starch combustion, generating smoke even under brief laser ablation. Thermal conductivity averaged 0.557 W/(mK) across all tests, consistent with prior work (Mohammadi et al., 2021). No statistically significant differences were observed between pre- and post-stimulation conditions (p>0.05). The phantom was fabricated and tested at room temperature using common, low-cost materials, highlighting its accessibility and versatility for multimodal stimulation studies (Mei et al., 2023; Linde et al., 2025; Saint-Martin & Avanaki, 2025).

### Limitations

Despite its versatility, the phantom had several limitations. Verification of its acoustic impedance against simulation data could be conducted through hydrophone testing (Fini & Tyler, 2020; Basso et al., 2024); however, this could not be performed. Residual water saturation within the phantom markedly reduced its overall electrical resistance, an effect which was spatially nonuniform across the structure (LaRocco et al., 2024). Thermal performance evaluation was restricted to surface measurements, as no internal sensors were employed to directly quantify thermal conductivity throughout the phantom (Farahani, 2019; Xu et al., 2010; Cho et al., 2023). While internal instrumentation would improve accuracy, finite element thermal modeling could provide valuable insight into spatial temperature gradients (Xu et al., 2010; Cho et al., 2023). The phantom also had a limited lifespan compared with abiotic analogs (Farahani, 2019; Kainz et al., 2019; Aboalgasm et al., 2024), and localized inconsistencies in gel–starch pocket formation reduced structural uniformity (Mei et al., 2023; Linde et al., 2025; Saint-Martin & Avanaki, 2025). Despite these limitations, its low fabrication cost and short production time enable high design flexibility and customization.

### Future work

Future work should focus on further optimizing the phantom for specific stimulation paradigms and validation across additional neuromodulation modalities. Although the current design was evaluated using LIFU and tDCS, it would be valuable to assess its performance under other widely used techniques such as DBS and TMS (Farahani, 2019; Xu et al., 2010; Cho et al., 2023). Optical characterization could also be conducted to refine its suitability for photobiomodulation applications and enable accurate photoacoustic imaging (Treeby et al., 2010). Enhancing structural complexity presents another avenue for advancement; this could include extending the 3D printed shell to incorporate features such as integrated pumping systems to simulate circulatory dynamics, and embedded sensors for real-time monitoring of temperature and heat distribution. The replacement of perishable components with durable materials would significantly increase the lifespan and reusability of the phantom (Mei et al., 2023; Linde et al., 2025; Saint-Martin & Avanaki, 2025). Development of a highly stable, long-lasting phantom would eliminate the need for fabrication prior to each test case and expand applicability in iterative and longitudinal studies (Cho et al., 2023; Mei et al., 2023; LaRocco et al., 2024; Linde et al., 2025; Saint-Martin & Avanaki, 2025).

## Conclusions

Measured values obtained using the fabricated phantom were consistent with multiple properties reported in the literature. Through integrating established concepts from acoustic, electrical, and thermal phantom designs, a low-cost, multimodal platform was produced and validated (Xu et al., 2010; Farahani, 2019; Cho et al., 2023; Mei et al., 2023; Linde et al., 2025; Saint-Martin & Avanaki, 2025). Simulations estimated a LIFU peak negative pressure of 0.631 MPa, closely matching the specified Pr.3 value, while producing a negligible temperature rise (<0.01 °C). Consistent with previous findings, tDCS did not induce lasting changes in the phantom’s material properties (LaRocco et al., 2024). Electrical conductivity was measured at 0.11±0.02 S/m, remaining relatively low due to water saturation within the phantom matrix (Mei et al., 2023; Linde et al., 2025; Saint-Martin & Avanaki, 2025). Thermal conductivity averaged 0.557 W/(mK), aligning with published values for brain-mimicking materials (Mohammadi et al., 2021). While evaluation focused on LIFU and tDCS, future work should extend characterization to other neuromodulation modalities such as DBS and TMS (Xu et al., 2010; Farahani, 2019; Cho et al., 2023). Optical property assessment would enhance applicability for photobiomodulation studies and enable accurate photoacoustic imaging (Treeby et al., 2010). Overall, fabrication of the phantom using common, inexpensive materials at room temperature underscores its accessibility and versatility as a platform for multimodal neuromodulation parameter optimization (Mei et al., 2023; Linde et al., 2025; Saint-Martin & Avanaki, 2025).

## Acknowledgments

I thank the Center for Neuroimaging, Neurophenotyping, Neurocomputation, and Neuromodulation (C4N) at The Ohio State University Wexner Medical Center for providing laboratory space for this research. I thank Joshua Gagliari of The Ohio State University College of Nursing for assisting with the brain mold.

## Supplementary information

The software and data is available at this site: https://github.com/psiwex/ihyBrainPhantom

